# Mapping Loci that Control Tuber and Foliar Symptoms Caused by PVY in Autotetraploid Potato (*Solanum tuberosum* L.)

**DOI:** 10.1101/156539

**Authors:** Washington da Silva, Jason Ingram, Christine A. Hackett, Joseph J. Coombs, David Douches, Glenn Bryan, Walter De Jong, Stewart Gray

**Affiliations:** School of Integrative Plant Science, Plant Pathology and Plant-Microbe Biology Section, Cornell University, Ithaca, NY 14853, U.S.A.; Biomathematics and Statistics Scotland, Invergowrie, Dundee, DD2 5DA, UK; Department of Plant, Soil and Microbial Sciences, Michigan State University, East Lansing, Michigan 48824; The James Hutton Institute, Invergowrie, Dundee, DD2 5DA, United Kingdom; School of Integrative Plant Science, Plant Breeding and Genetics Section, Cornell University, Ithaca, NY 14853, U.S.A.; USDA, ARS, Emerging Pests & Pathogens Research Unit, Ithaca, NY 14853, U.S.A.

**Keywords:** genetic linkage map, QTL, autotetraploid potato, single-nucleotide polymorphism, *Potato Virus Y*, PTNRD

## Abstract

Potato tuber necrotic ringspot disease (PTNRD) is a tuber deformity associated with infection by the tuber necrotic strain of *Potato virus Y* (PVY^NTN^). PTNRD negatively impacts tuber quality and marketability and poses a serious threat to seed and commercial potato production worldwide. PVY^NTN^ symptoms differ in the cultivars Waneta and Pike: Waneta expresses severe PTNRD and foliar mosaic with vein and leaf necrosis, whereas Pike does not express PTNRD and mosaic is the only foliar symptom. To map loci that influence tuber and foliar symptoms, 236 F_1_ progeny of a cross between Waneta and Pike were inoculated with PVY^NTN^ isolate NY090029 and genotyped using 12,808 Potato SNPs. Foliar symptom type and severity were monitored for 10 weeks, while tubers were evaluated for PTNRD expression at harvest and again after 60 days in storage. Pairwise correlation analyses indicate a strong association between PTNRD and vein necrosis (τ = 0.4195). QTL analyses revealed major-effect QTLs on chromosomes 4 and 5 for mosaic, 4 for PTNRD, and 5 for foliar-necrosis symptoms. Locating QTLs associated with PVY-related symptoms provides a foundation for breeders to develop markers that can be used to screen out potato clones with undesirable phenotypes, e.g., those likely to develop PTNRD or to be symptomless carriers of PVY.

## INTRODUCTION

*Potato virus Y* (PVY) is the most common and most serious virus affecting US potato production, and resistant potato cultivars represent the most effective control option (Karasev and Gray 2013a; Fulladolsa et al. 2015). PVY exists as a myriad of strains, including: the ordinary strain PVY^O^, the tobacco vein necrosis strain PVY^N^, the stipple streak strain PVY^C^, and the tuber necrosis strain PVY^NTN^ that elicits potato tuber necrotic ringspot disease (PTNRD) (Karasev and Gray 2013a; Schubert et al. 2007). Until recently, North American potato breeding programs have not prioritized PVY resistance during selection. A lack of resistance and the popularity of several widely-planted varieties that are symptomless carriers of PVY have facilitated an increase in PVY incidence and contributed to the emergence of new PVY strains that cause PTNRD (Gray et al. 2010; Karasev and Gray 2013b). PTNRD poses a serious threat to the seed and commercial production industries by contributing to the rejection of seed lots for exceeding virus tolerance, as well as negatively impacting tuber quality (Karasev and Gray 2013a; Kerlan and Moury 2008). Some potato cultivars widely grown in the US and Canada are highly susceptible to PTNRD, such as Yukon Gold, Yukon Gem, Red Norland, Highland Russet, Alturas, Blazer, and Ranger Russet (McDonald and Singh 1996; Singh et al. 1998).

Resistance genes effective against PVY have been identified in cultivated and wild potato species (Cockerham 1970; Jones 1990; Fulladolsa et al. 2015; Karasev and Gray 2013a) and have been classified into two types, hypersensitive resistance (HR) and extreme resistance (ER) (Gebhardt and Valkonen 2001). HR is associated with the development of visible necrotic lesions at the point of infection. In some varieties the response can be a systemic necrosis manifested as vein necrosis, leaf necrosis or leaf drop. All of these responses can contribute to limiting virus replication and systemic spread, as well as reducing aphid transmission efficiency of the virus from these plants. HR is conferred by *N* genes (Solomon-Blackburn and Barker 2001). The major *N* genes, *Ny*_*tbr*_ and *Nc*_*spl*_ (Celebi-Toprak et al. 2002; Moury et al. 2011), *Ny-1* (Szajko et al. 2008), and *Ny-2* (Szajko et al. 2014) have been mapped to chromosomes 4, 9, and 11, respectively. ER is asymptomatic, results in no detectable virus multiplication in inoculated plants, and is conferred by *R* genes (Solomon-Blackburn and Barker 2001). Several molecular markers have been developed for potato *R* genes, including: RYSC3 for detection of *Ry*_*adg*_ from *S. tuberosum* ssp. *andigena*, on chromosome 11 (Sorri et al. 1999; Kasai et al. 2000); 38–530 and CT220 for *Ry*_*chc*_ from *S. chacoense*, on chromosome 9 (Hosaka et al. 2001; Sato et al. 2006); and GP122, STM003, and YES3-3B for *Ry*_*sto*_ from *S. stoloniferum*, on chromosome 12 (Song et al. 2005; Song and Schwarzfischer 2008; Valkonen et al. 2008). Many of those markers have been successfully incorporated in breeding programs to develop PVY–resistant cultivars (Fulladolsa et al. 2015; Ottoman et al. 2009; Watanabe 2015).

Marker-assisted selection (MAS) has proven to be a fast and efficient tool to select cultivars with desirable traits in plant breeding (Xu and Crouch 2008). Developing markers linked to important genes in cultivated potato (*Solanum tuberosum* ssp. *tuberosum*) is more challenging than in many other crops, primarily because conducting linkage analyses is more difficult in autotetraploids than in diploids. Nevertheless, with the sequencing of the potato genome (PGSC 2011), followed by the development, validation, and release of the Infinium Potato SNP Arrays (Hamilton et al. 2011; Felcher et al. 2012), improvements of statistical models for analyzing SNP dosage in tetraploids (Hackett et al. 2013; Hackett et al. 2014; Preedy and Hackett 2016; Hackett et al. 2001), and the development of TetraploidSNPMap user-friendly software specifically designed to analyze SNP markers in polyploid germplasm (Hackett et al. 2017) – QTL analyses in potato have recently become much more feasible.

Developing varieties that do not express PTNRD upon infection is potentially a useful complement or alternative to developing varieties resistant to PVY. Genetic markers that breeders could use to select for lack of PTNRD expression would facilitate the development of such varieties. The goal of this research was to map genes that mediate PTNRD and other types of foliar symptoms induced by PVY infection (mosaic, vein necrosis, and leaf necrosis).

## MATERIALS AND METHODS

### Plant Material

The H25 mapping population comprises 236 F_1_ progeny of a cross between the cultivars Waneta (as female) and Pike (as male). These two cultivars express different symptoms when infected by PVY isolate NY090029 (a PVY^NTN^ strain). Waneta expresses severe PTNRD and foliar mosaic with vein and leaf necrosis. Pike does not exhibit PTNRD and mild mosaic is the only foliar symptom.

True potato seeds of H25 were germinated on a bed of Cornell potting mix (Boodley and Sheldrake 1982). After one month, 80 seedlings were individually transplanted to 15-cm clay pots. Each seedling was vegetatively propagated via cuttings to increase the number of plants per genotype. One tuber of each parent was individually planted in a 15-cm clay pot and cuttings were taken from the sprouts, also to increase the number of plants per genotype. All cuttings, from parents and progeny, were dipped in Hormex rooting hormone #1 (Brooker Chem. Corp., Chatsworth, CA) and planted individually into 96 well trays containing Cornell soil mix for rooting and grown for one month. Additionally, 200 true potato seeds from the H25 population were sterilized and placed into tissue culture media by the following method. Seeds were soaked overnight in a 1500ppm Gibberellic acid solution, then the solution was removed and a 10% bleach solution was added and incubated for 10 minutes with periodic inverting of the tube. The bleach solution was removed and sterile H_2_O was added to wash the seeds, repeating the washing step four times. Seeds were plated onto a sterile autoclaved size 1 Whatman circle filter paper in a petri dish damped with H_2_O. Petri dishes were sealed with parafilm and placed under growth lights (16hr light/day) until the seeds sprouted. The young sprouts were then transferred to Murashige and Skoog medium. 156 progeny were established in tissue culture and these plants, as well as the progeny sown directly into soil, were used for virus phenotyping and SNP genotyping (“the mapping population”). Six well-rooted plants from each clone of the mapping population and the parents were transplanted individually in four-liter plastic pots containing Cornell soil mix. During all steps of the experiment, including germination and sprouting, plants were maintained in an insect-free greenhouse under 16h days at 25±3°C.

### Phenotypic data

Two weeks after transplanting, five plants from each clone and the two parents were inoculated with PVY^NTN^ isolate NY090029. One plant from each clone and each parent was left uninoculated as a negative control. To prepare viral inoculum, the PVY^NTN^ isolate NY090029 (maintained in lyophilized tobacco tissue at −80°C) was mechanically inoculated to individual tobacco (*Nicotiana tabacum*) plants at the three-to five-leaf stage. Lyophilized tissue (100 mg) was homogenized in 500 µl of phosphate-buffered saline (PBS) (137 mM NaCl, 2.7 mM KCl, 10 mM Na_2_HPO_4_, 2 mM KH_2_PO_4_, and pH adjusted to 7.4 with HCl) and rubbed onto carborundum-dusted (325 mesh) tobacco leaves. Inoculated plants were kept in a greenhouse for one month. Infected tobacco leaves were harvested, ground in PBS in a volume of 1:5 (1g of leaf to 5ml of PBS), and filtered through cheesecloth to produce the inoculum. Then, potato plants were inoculated by using a cotton swab to lightly rub the PVY inoculum using carborundum as an abrasive. The same plants were inoculated twice more, with one-week intervals between inoculations. Plant infection status was checked by ELISA using a 4C3 commercial kit (Agdia Inc., Elkart IN, USA), following the manufacturer’s directions.

One and three weeks after the final round of inoculations, foliar symptoms were evaluated on each plant (Figure 1). Mosaic was scored on this scale: 0 = no symptoms, 1= mild mosaic (mosaic pattern muted but present), 2 = typical mosaic (mosaic pattern evident and some leaf rugosity possible), and 3 = severe mosaic (mosaic pattern evident, plant stunting, rugosity and deformation on leaves). The grading for leaf and vein necrosis symptoms was binary: 0 = no symptoms, 1 = symptoms present. Four months after the potato cuttings were transplanted, the vines were removed, the pots were left to dry out for three weeks, and then tubers were harvested. At harvest and three months later, PTNRD severity was visually evaluated for each tuber as follows: 0= no PTNRD, 1= 1-10% PTNRD, 2= 11-25% PTNRD, 3= 26-50% PTNRD, 4= 51-75% PTNRD, and 5= 76-100% PTNRD (Figure 2). For subsequent analyses the highest disease value – the most severe symptoms observed among the five plants tested for each genotype – was used. In a pilot study, using the linkage maps from the full population (236 clones), we ran QTL analyses on a subset of the population (85 clones) using the mean and the highest disease values, and found the same significant QTLs for both types of data; we elected to use the highest disease values in all subsequent analyses. Pairwise correlation analyses were performed on the phenotypic dataset with the non-parametric Kendall’s tau rank correlation coefficient to measure the strength of the relationship between each type of symptom. All statistical analyses and plotting for data visualization were performed in R (R Core Team 2016) using the R packages Hmisc (version 4.0-0) (Harrell Jr. 2016) and corrplot (version 0.77) (Wei and Simko 2016).

**Figure 1.**
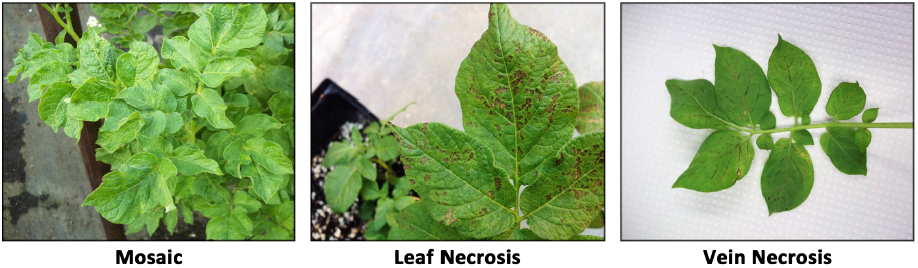
Foliar symptom severity ratings for 236 clones from the H25 population. Ratings were scored on a 0 to 3 scale with 0 = no disease and 3 = most severe symptoms for mosaic and a 0 to 1 scale for leaf and vein necrosis with 0 = no disease and 1 = disease.

**Figure 2.**
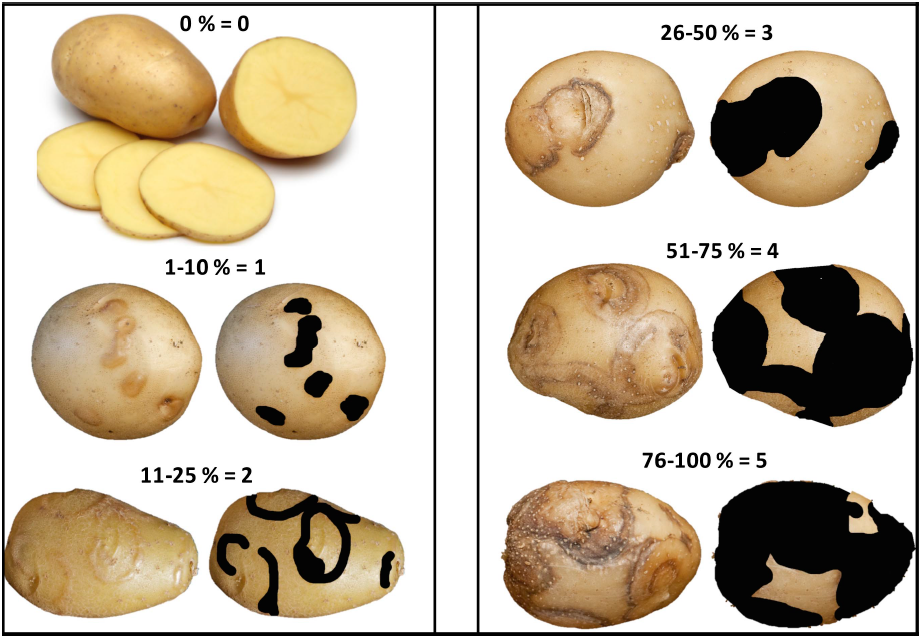
PTNRD severity rating of the tubers from 236 clones of the H25 population. Ratings were based on a 0 to 5 scale with 0 = no disease and 5 = most severe symptoms.

### SNP genotyping

DNA from 236 progeny clones and their parents was extracted from frozen-leaf-tissue using a QIAGEN DNeasy Plant Mini Kit (Qiagen, Valencia, CA-USA), following the manufacturer’s directions. DNA was quantified with the Quant-it PicoGreen assay (Invitrogen, San Diego, CA-USA) and adjusted to a concentration of 50ng µL^−1^. The population was genotyped with the Illumina Infinium V2 Potato SNP Array (12,808 SNPs: original SolCAP Infinium 8303 Potato SNP Array with 4,500 additional SNPs to increase coverage in candidate genes and R-gene hotspots) (Hamilton et al. 2011). Illumina GenomeStudio software (Illumina, Inc., San Diego, CA-USA) was used for initial sample quality assessment and generating marker theta values (which give dosage allelic information for parents and offspring). In an autotetraploid mapping population, five allele dosages (AAAA, AAAB, AABB, ABBB, and BBBB) are possible and are expected to consist of theta scores in five clusters, centering around 0.0, 0.25, 0.50, 0.75, and 1.0, respectively. Tetraploid (5-cluster) genotyping was based on theta value thresholds, using a custom script from the SolCAP project (Hirsch et al. 2013). Using this script, 5-cluster calling and filtering were performed to remove low quality markers and markers with multiple hits to the potato genome sequence of *Solanum tuberosum* group Phureja DMI-3 516 R44 (Sharma et al. 2013). SNPs with >20% missing genotype calls in the population were excluded from the dataset.

### Linkage map construction and QTL analysis

Construction of linkage maps and QTL analysis of each chromosome were performed as described in (Hackett et al. 2014; Hackett et al. 2013; Preedy and Hackett 2016; Hackett et al. 2017). All linkage and QTL analyses involving testing for distorted segregation, clustering analysis, calculation of recombination fractions and LOD (logarithm of the odds) scores, ordering of SNPs, and inference of parental phase, were performed in TetraploidSNPMap. Markers with significance of the χ^2^ goodness-of-fit statistic less than 0.001 for simplex SNPs and 0.01 for duplex or greater dosage SNPs were flagged as distorted. To detect and remove problematic markers and for ordering of SNPs, the following analyses were performed: hierarchical clustering analyses using average linkage clustering of SNPs with expected ratios, 2-point analyses to calculate the recombination frequency and LOD score for the SNPs pairs in each possible phase, and multidimensional scaling analysis (MDS) to calculate the best order for the SNPs in the linkage group (Preedy and Hackett 2016; Hackett et al. 2017). Finally, the phases of the ordered SNPs were inferred as far as possible by the automated phase analysis in TetraploidSNPMap and completed manually prior to carrying out QTL analysis.

QTL analysis was run for each linkage group separately using three input files: the linkage map, the SNP data for the linkage group, and the phenotypic trait dataset. For each trait, interval mapping displayed the LOD profile on the chromosome, giving the LOD score statistics, percentage variation explained, and QTL effect for each homologous chromosome. 90% and 95% LOD thresholds were obtained to establish the statistical significance of each QTL position using permutation tests with 300 permutations. Simple models for the genotype means estimated at the most probable QTL position were calculated using the Schwarz Information Criterion (SIC) (Schwarz 1978), models with the lowest value for SIC are considered the best models (Hackett et al. 2014). Linkage maps and QTL positions were generated in MapChart 2.30 (Voorrips 2002).

Concordance between the linkage maps generated in this study and the potato reference genome (PGSC Version 4.03 Pseudomolecules) was evaluated in MareyMap R package version 1.3.1 (Rezvoy et al. 2007). Plots of the genetic position (cM) with the physical position (Mb) of each SNP marker in each chromosome were generated using the graphical interface MareyMapGUI, the interpolation method “cubic splines” was used to calculate the curve slope.

### Data availability

All the raw data from this study was compiled in txt tables and are available in the supplemental files: Table S1 and Table S2. Complementary information for the Results and Discussion section are provided in Support Information: Figures S1, S2, and S3, and Tables S3 and S4.

## RESULTS AND DISCUSSION

### Genotyping and preliminary SNP marker processing

The 12808 SNPs from the new Illumina Potato V2 SNP Array (12K) were used to genotype the parents and 236 offspring in this study. After a pre-filtering step to remove SNPs with missing theta values, low quality, and those with multiple hits to the potato reference genome PGSC Version 4.03 Pseudomolecules, 4,859 SNPs were selected for downstream analyses (Supporting Information, Table S1). Of these, 1063 SNPs had missing data in >20% of the population, and were also excluded from the dataset. The remaining 3796 SNPs were loaded into TetraploidSNPMap and 1258 distorted SNPs with chi-square statistics having a significance less than 0.001 were removed. Hierarchical clustering analyses easily grouped the remaining 2538 markers into 12 linkage groups (Supporting Information, Table S2). A total of 95 SNPs was flagged as duplicated and 17 were excluded as outliers after clustering, 2-point, and MDS analyses.

Approximately, 65% (1583) of the markers followed the parental genotype configurations of simplex (AAAA X AAAB, AAAB X AAAA), duplex (AAAA X AABB, AABB X AAAA), and double-simplex (AAAB X AAAB, ABBB X ABBB), while ~ 35% (843) were between simplex-duplex (AAAB X AABB) and double-duplex (AABB X AABB) configurations (Table S3). The large number and diversity of configurations of SNPs in our dataset allowed for the construction of high-density linkage maps, which significantly increased the chances for the detection of significant QTLs for the traits studied (Massa et al. 2015; Hackett et al. 2013; Li et al. 2014; Hackett et al. 2014).

### Linkage map construction and QTL analysis

The 2426 SNPs were mapped to the 12 potato chromosomes with chromosomes 1 and 12 having the highest and the lowest number of mapped SNPs (281 and 138), respectively (Table 1). Overall, 1809 SNPs segregated in “Waneta”, 1962 segregated in “Pike”, and 1345 SNPs segregated in both parents. The total genetic distance for each of the parental maps was 1052.6 cM (for Waneta) and 1097.1 cM (for Pike), with the map lengths of individual chromosomes ranging from 72.1 to 120.6 cM. There was an average of 157 SNP markers per chromosome and a marker density of ~1.75 SNPs per cM. The genetic maps of both parents, covered, on average, 98% of the PGSC v4.03 Pseudomolecules (Table 1, and Figure 3).

**Table 1.**
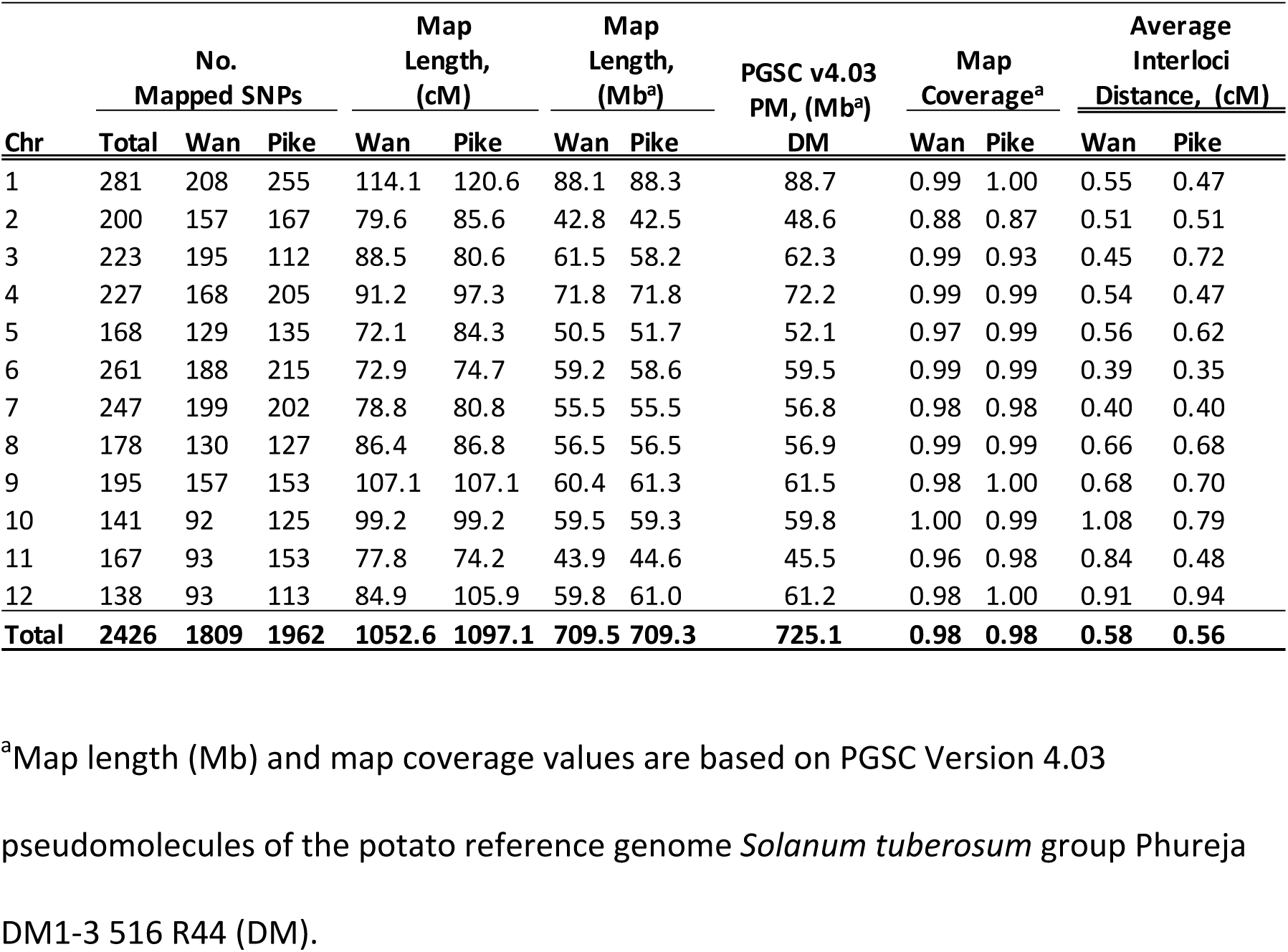
Summary of the parental linkage maps, Waneta (Wan) and Pike.

**Figure 3.**
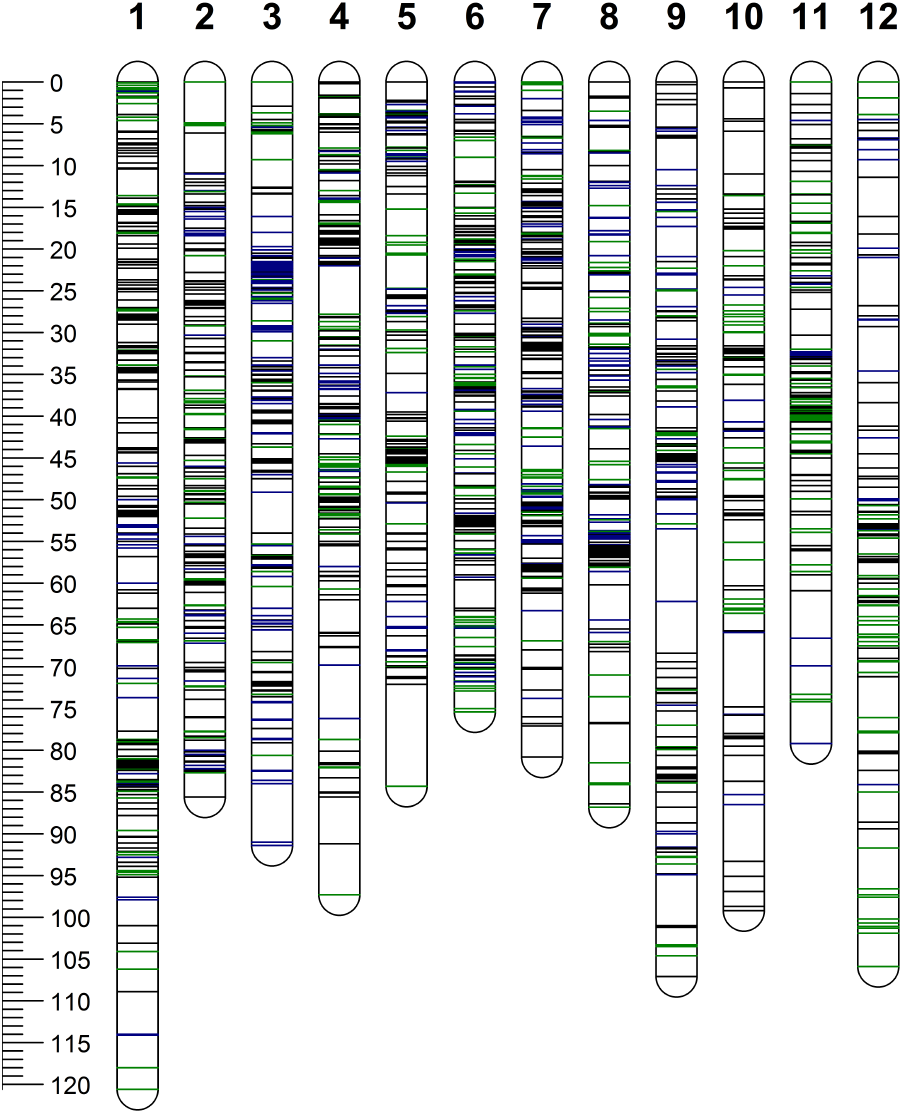
Distribution of single-nucleotide polymorphism (SNP) markers on 12 chromosomes (1-12) of the parents (Waneta and Pike). The scale bar shows the genetic distance in cM. SNPs positions are represented by green lines (Waneta), blue lines (Pike), and black lines (both parents) across each chromosome.

### Vein Necrosis positively correlated with PTNRD

Non-parametric Kendall’s tau rank correlation analyses indicated a weak correlation among mosaic and other symptom types (PTNRD, foliar necrosis, and vein necrosis). In contrast, vein necrosis exhibited the highest correlation with other symptom types especially PTNRD (Figure 4) – an indication that when vein necrosis is observed, there is a high chance of PTNRD development in tubers. The evaluation of PTNRD requires a lot of time as tubers need to be stored for at least two months after harvest for full expression of the symptoms. Knowing that vein necrosis is correlated with PTNRD may benefit potato growers and researchers alike.

**Figure 4.**
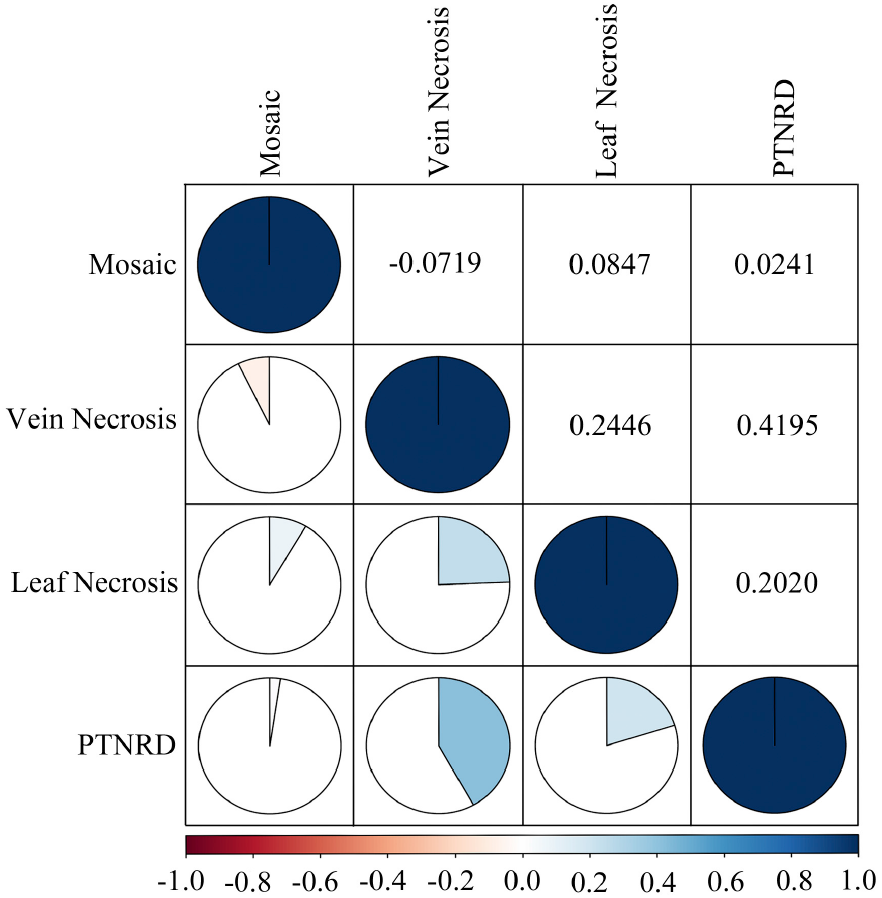
Pairwise correlation analyses using the non-parametric Kendall’s tau rank correlation coefficient to measure the strength of the relationship between each type of symptom expression. Positive correlations are displayed in blue and negative correlations in red. Color intensity is proportional to the correlation coefficients.

### Significant QTLs were identified on chromosomes 4 and 5 for mosaic and leaf necrosis

Mosaic symptoms were frequent in the population, with 219 of the 236 offspring expressing symptoms (Figure 5). This was not surprising, as we had found in preliminary studies that PVY isolate NY090029 is highly virulent and elicited severe mosaic in most inoculated plants including both parents. In contrast, only 31 and 172 clones developed leaf necrosis and vein necrosis, respectively (Figure 5). QTL analyses revealed significant QTLs on chromosomes 4 and 5 for mosaic and leaf necrosis (Table 2 and Figures 6, 7, S1, S2). No significant QTLs were detected for vein necrosis in the population.

**Table 2.**
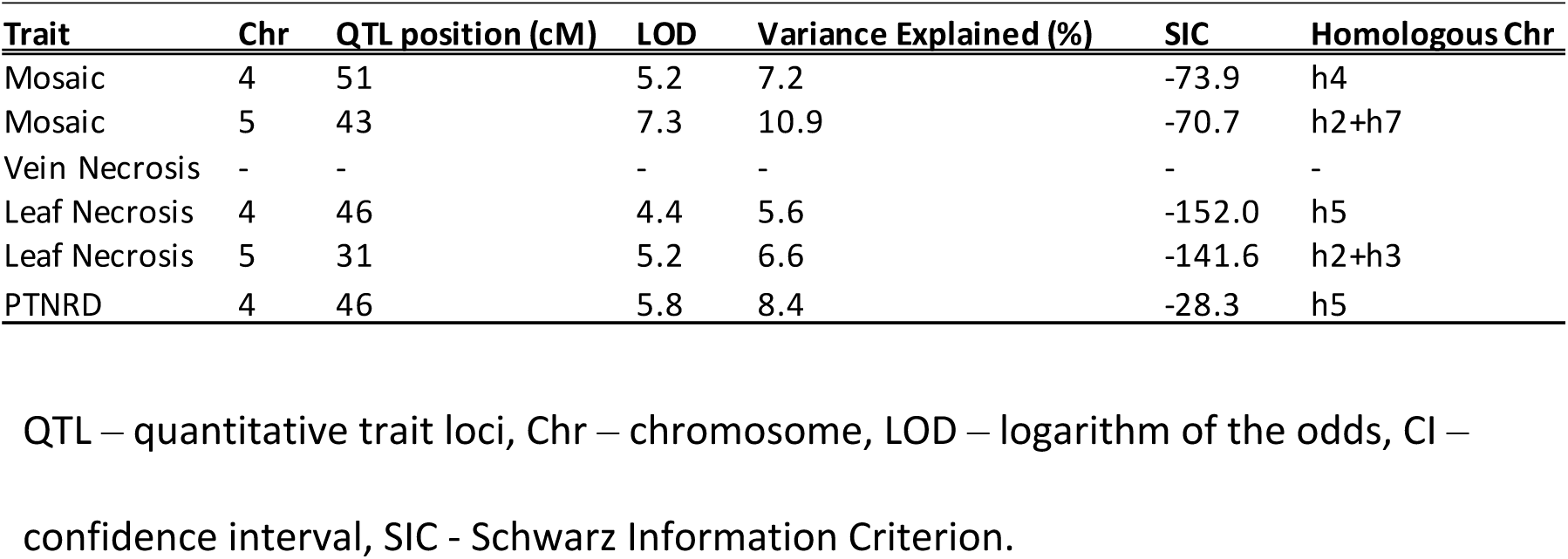
QTL information for the traits analyzed in the H25 population. QTL – quantitative trait loci, Chr – chromosome, LOD – logarithm of the odds, CI –confidence interval, SIC - Schwarz Information Criterion.

| Trait | Chr | QTL position (cM) | LOD | Variance Explained (%) | SIC | Homologous Chr |
| --- | --- | --- | --- | --- | --- | --- |
| Mosaic | 4 | 51 | 5.2 | 7.2 | -73.9 | h4 |
| Mosaic | 5 | 43 | 7.3 | 10.9 | -70.7 | h2+h7 |
| Vein Necrosis | - | - | - | - | - | - |
| Leaf Necrosis | 4 | 46 | 4.4 | 5.6 | -152.0 | h5 |
| Leaf Necrosis | 5 | 31 | 5.2 | 6.6 | -141.6 | h2+h3 |
| PTNRD | 4 | 46 | 5.8 | 8.4 | -28.3 | h5 |

**Figure 5.**
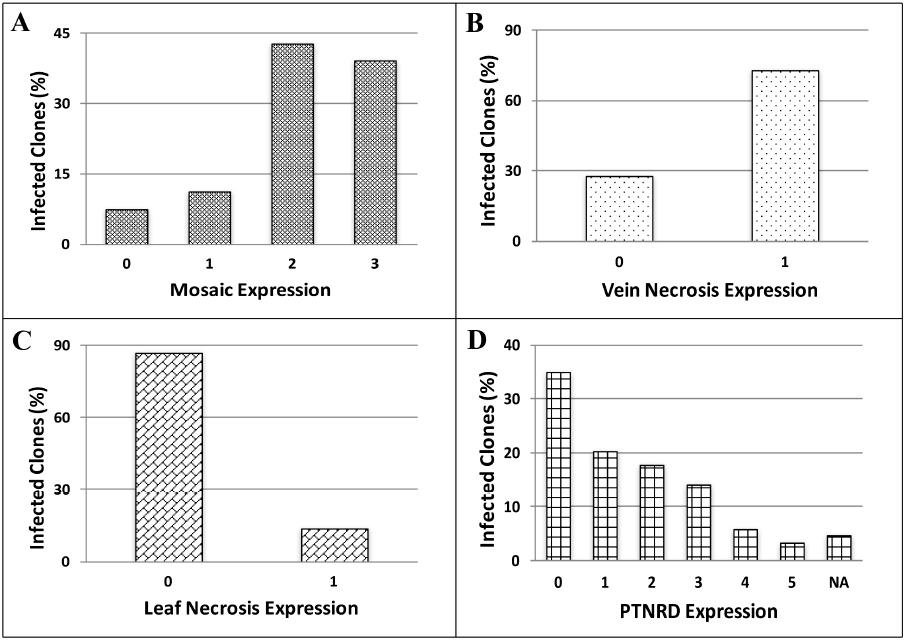
Percentage of Waneta x Pike offspring expressing varying degrees of foliar (A) Mosaic, (B) Vein Necrosis, (C) Leaf Necrosis, and tuber symptoms (D) PTNRD. See Figures 1 and 2 for symptom scales. NA = number of clones that did not produce tubers; PTNRD was not evaluated with them.

**Figure 6.**
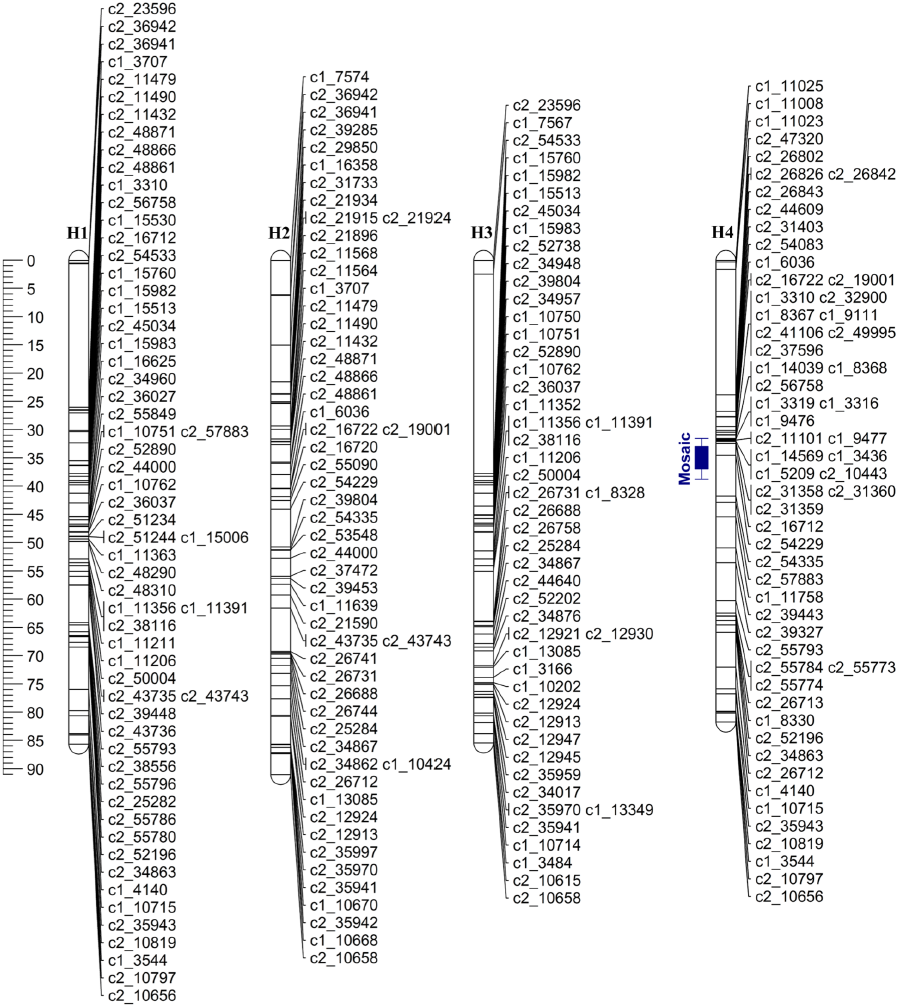
Linkage map of Waneta chromosome 4 (H1-H4 = homologous maps). The blue bar corresponds to the 95% support LOD interval for the QTLs locations for leaf necrosis and mosaic, respectively. Whiskers represent the two LOD support interval and the solid box represents the one LOD support interval for the QTL location.

**Figure 7.**
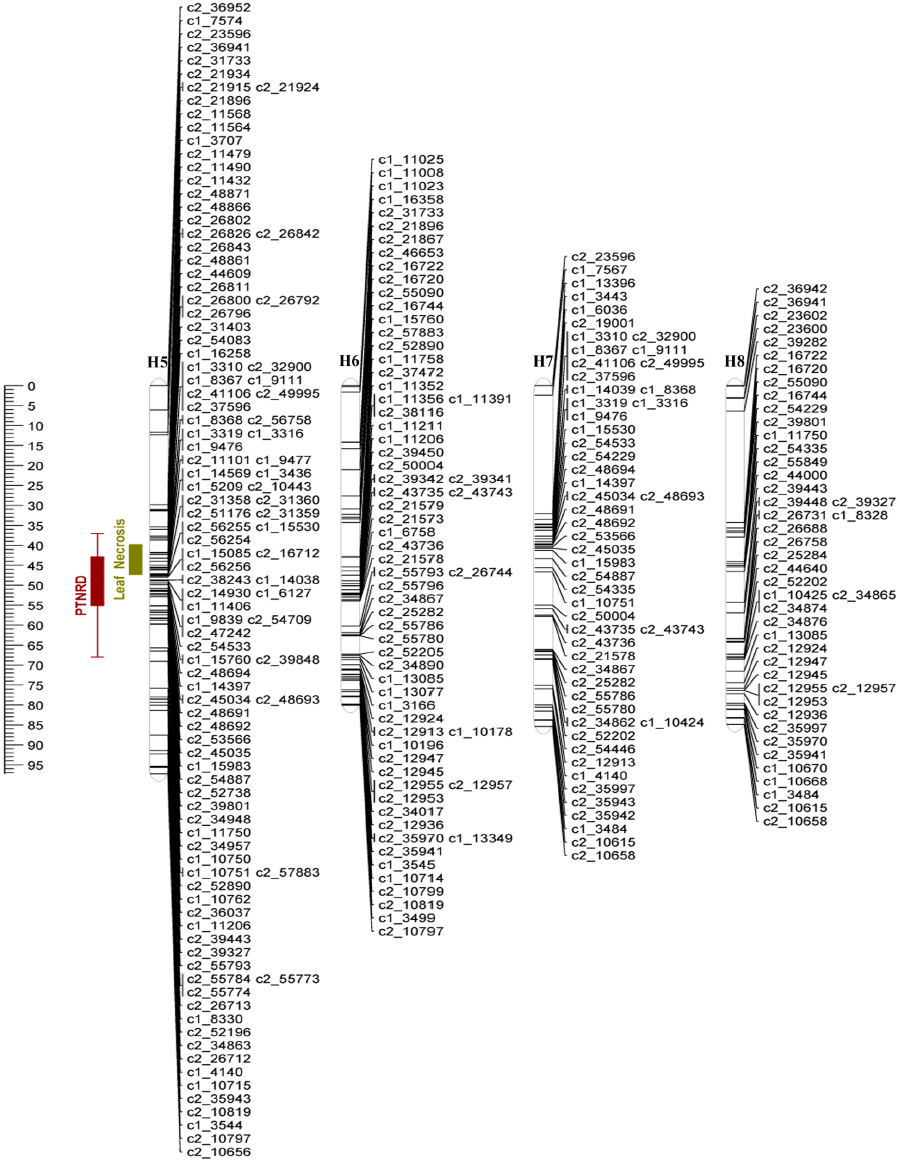
Linkage map of Pike chromosome 4 (H5-H8 = homologous maps). The brown and olive green bars correspond to the support LOD intervals for the QTLs location for PTNRD and Leaf Necrosis, respectively. For each bar, whiskers represent the two LOD support interval and the solid box represents the one LOD support interval for the QTL location.

On chromosome 4, the QTLs had maximum LOD scores of 5.20 and 4.44 explaining 7.2% and 5.6% of the trait variances for mosaic and leaf necrosis, respectively. These LOD scores were above the upper 95% LOD permutation thresholds of 3.95 and 3.81 and the QTL peaks were located at positions 51 cM and 46 cM for mosaic and leaf necrosis, respectively. Analyses of different simple genetic models were performed with TetraploidSNPMap to determine the best simple fitting model for each trait. For mosaic, the best model was a simplex allele (AAAB) on homologous chromosome 4 (H4, Fig. 6) of Waneta, with the B allele associated with a decrease in symptom expression. This model had the lowest SIC, −73.94, in comparison with the full model (SIC = −55.94). For leaf necrosis, the best model was a simplex allele (BAAA) on homologous chromosome 5 of Pike (H5, Fig. 7), with the B allele associated with a decrease in symptom expression. This model had SIC = −151.98, while the SIC for the full model was −145.80.

On chromosome 5, the maximum LOD scores were 7.34 and 5.20 and those QTLs explained 10.9% and 6.6% of the phenotypic variance for mosaic and leaf necrosis, respectively (Table 2). The LOD peaks were located at positions 43 cM and 31 cM and their scores were above the upper 95% LOD permutation thresholds of 3.65 and 3.76 for mosaic and leaf necrosis, respectively. The simpler models analyses estimated a double-simplex and a duplex genotype for mosaic and leaf necrosis, respectively. For mosaic, the best model was an ABAA X AABA configuration on homologous chromosomes 2 and 7 (H2+H7, Figures S1 and S2) with the B allele associated with a decrease in symptom expression, and both parents contributing the B allele to their offspring. This model had the lowest SIC, −70.68, in comparison with the full model (SIC = −66.20). For leaf necrosis, the best model was an ABBA configuration on homologous chromosomes 2 and 3 of Waneta (H2+H3, Figure S1) with the B allele associated with a decrease in symptom expression. The SIC for this model was −141.62, the full model had SIC = −136.41.

Analyses of the concordance between the linkage maps and the potato reference genome (PGSC Version 4.03 Pseudomolecules) for chromosomes 4 and 5 generated graphs that were consistent with published chromosome structures (Figure S3) (Massa et al. 2015; Felcher et al. 2012; Sharma et al. 2013).

### A major-effect QTL for PTNRD expression was detected on chromosome 4

One hundred and forty-five clones produced tubers that expressed some degree of PTNRD. Of the 89 remaining clones, 11 clones did not produce tubers and 78 produced tubers with no PTNRD. A PTNRD QTL was detected on chromosome 4 that had a LOD score of 5.82, explained 8.6% of the trait variance (Table 2), and was above the 95% LOD permutation upper threshold of 3.92. The QTL peak was located at 46 cM and analyses of different genetic models indicated that an allele from Pike explains the trait variance. The QTL is linked to a simplex SNP (AAAA x BAAA), with the B allele associated with a decrease in disease on homologous chromosome 5 (H5, Figure 7). This model had the lowest SIC of – 28.32 compared to the full model with SIC = −23.61. The closest SNP with this configuration is the SNP solcap_snp_c2_39848 at genetic position 47.09 cM and physical position 35.68 Mb. This QTL was located in the central region of chromosome 4, the same region where QTLs for mosaic and leaf necrosis were detected. The center of chromosome 4 harbors two known genes, N*y*_*tbr*_ and N*c*_*spl*_, that cause HR in potatoes when infected with PVY^O^ and PVY^C^, respectively (Celebi-Toprak et al. 2002; Moury et al. 2011). It is possible that alleles of these genes influence PTNRD, mosaic, and/or leaf necrosis symptoms. R genes frequently occur in tightly linked clusters (Michelmore and Meyers 1998) and the distribution of such genes and QTLs is not random in the potato genome (Gebhardt and Valkonen 2001). The detection of major QTLs for different PVY symptom types in close proximity to each other on chromosome 4 suggests that markers diagnostic for specific haplotypes of this region may prove useful for breeders who want to select genes that confer resistance to infection and/or multiple PVY-related symptoms. Finally, it is important to point out that QTL analysis is approximate, as the disease traits evaluated in this study are ordinal or binary scores and so definitely not normal. However, basing significance on permutation of this data helps, in part, to address this problem.

## ACKNOWLEDGEMENTS

The authors would like to thank Kelly Zarka and Natalie Kirkwyland for helping with tissue culture and Daniel Zarka for genotyping the H25 population. This study was funded in part by USDA-SCRI (2009-51181-05894 and 2014-51181-22373), the UK Biotechnology and Biological Sciences Research Council grant # BB/L011840/1 as part of the joint USDA-NSF-NIH-BBSRC Ecology and Evolution of Infectious Diseases program. The work of Glenn Bryan, Christine Hackett, the TetraploidSNPMap software development were funded by the Rural & Environment Science & Analytical Services Division of the Scottish Government. Washington da Silva was partially supported by a SUNY Diversity Fellowship and by the 2014/2015 National Potato Council (NPC) scholarship.

## Supplemental Figures

**Figure S1.**
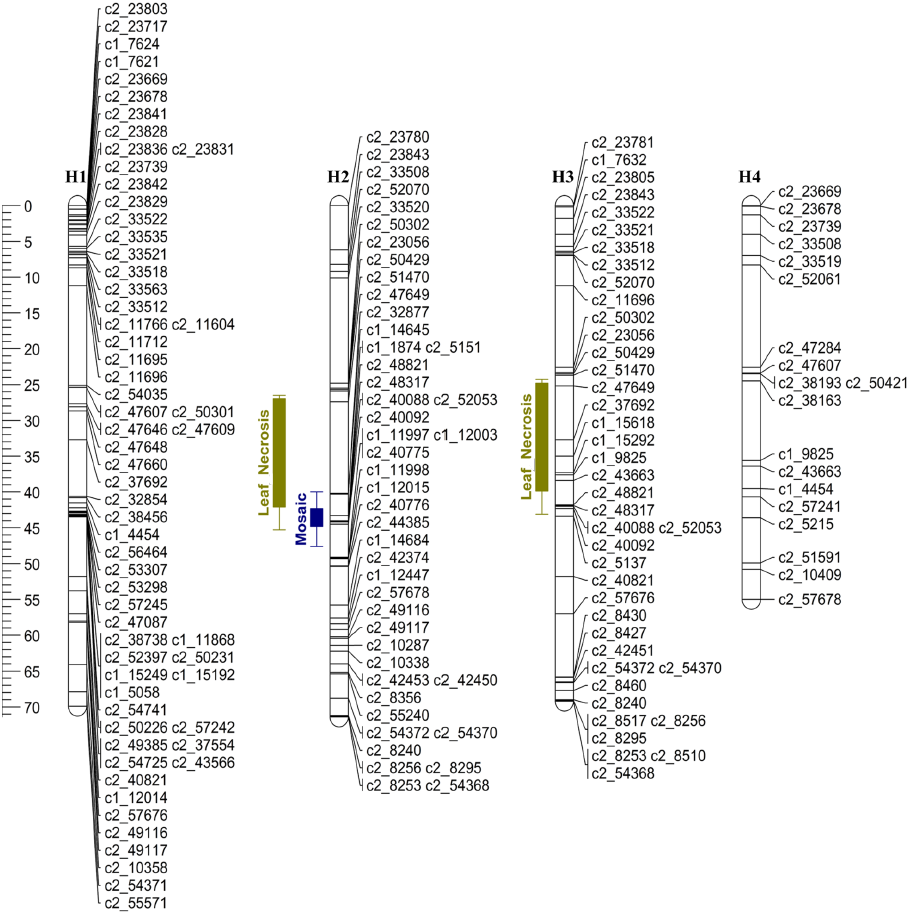
Linkage map of Waneta chromosome 5 (H1-H4 = homologous maps). The olive green and blue bars correspond to the support LOD interval for the QTLs location for leaf necrosis and mosaic symptoms, respectively. In the bars, whiskers represent the two LOD support interval and the solid box represents the one LOD support interval for the QTL location.

**Figure S2.**
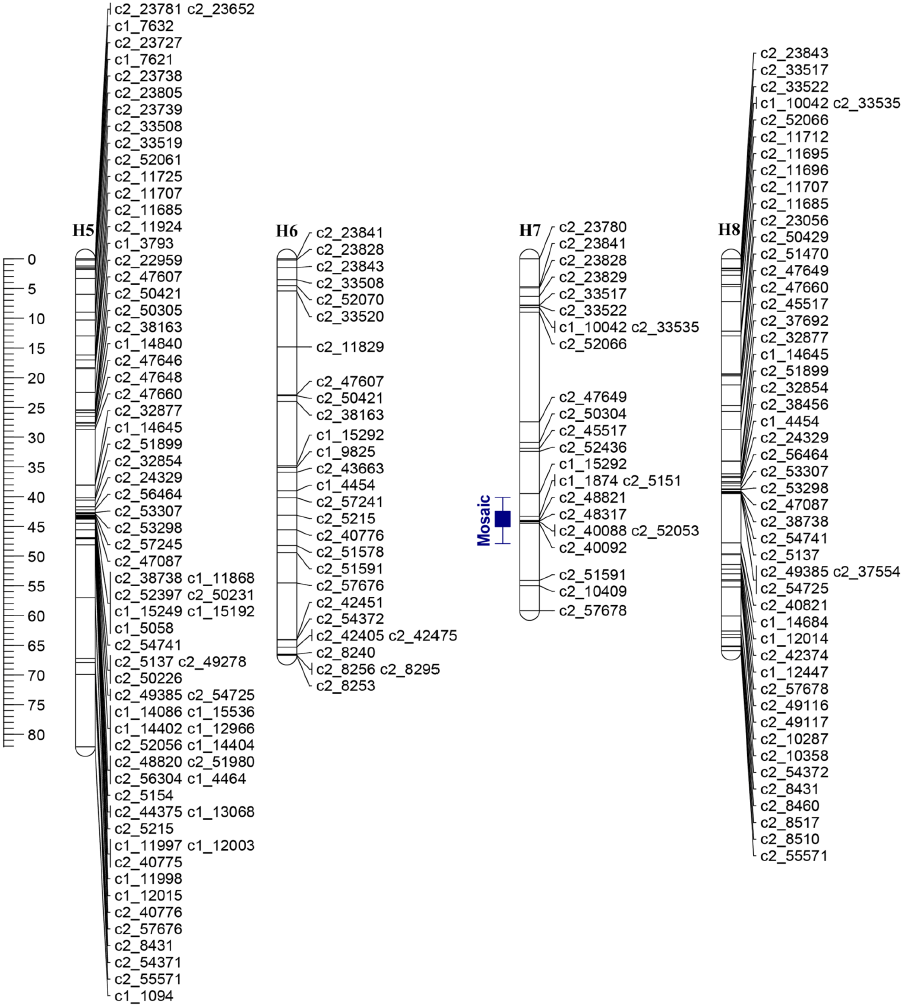
Linkage Map of Pike chromosome 5 (H5-H8 = homologous maps). The blue bar corresponds to the support LOD interval for the QTL location for Mosaic. In the bar, whiskers represent the two LOD support interval and the solid box represents the one LOD support interval for the QTL location.

**Figure S3.**
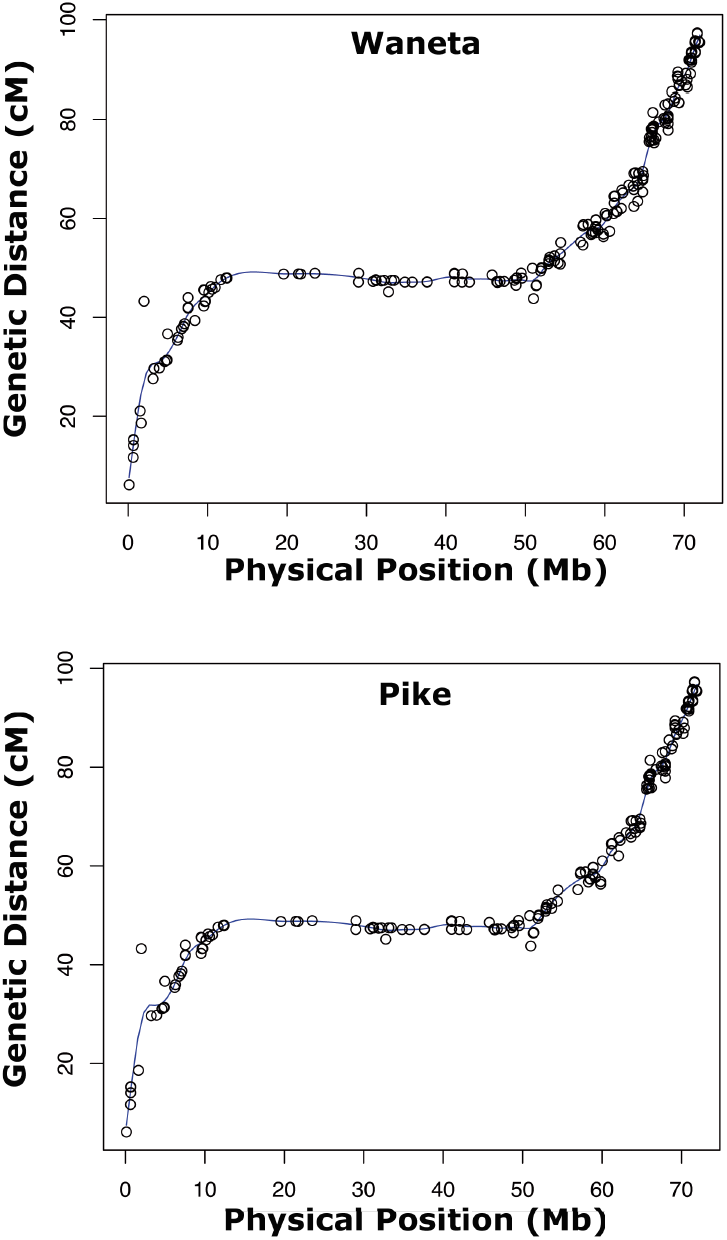
Graph of chromosome 4 from the parents (Waneta and Pike) showing the genetic location (cM) and the physical position (Mb) of SNP markers.

## Supplemental Tables

**Table S4.**
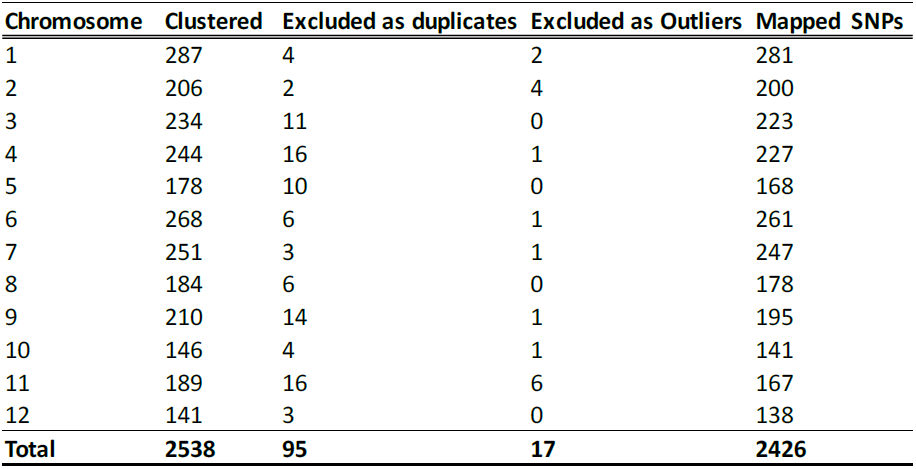
The number of SNPs clustered, excluded as duplicates and outliers, andmapped on each chromosome by TetraploidSNPMap Software.

**Table S3.**
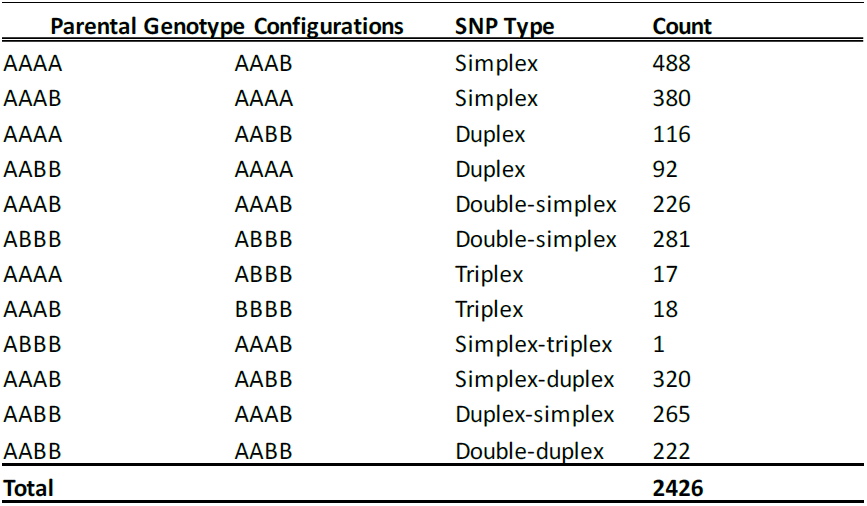
Parents and F1 offspring genotype configurations in the H25 tetraploid mapping population.

## LITERATURE CITED

Boodley, J.W., and R. Sheldrake, 1982 Cornell peat-like mixes for commercial plant growing. Cornell Inf. Bull. 43.

Celebi-Toprak, F., S.A. Slack, and M.M. Jahn, 2002 A new gene, Ny_tbr_, for hypersensitivity to Potato virus Y from Solanum tuberosum Maps to Chromosome IV. Theoretical and Applied Genetics 104 (4):669–674.

Cockerham, G., 1970 Genetical studies on resistance to potato viruses X and Y. Heredity 25 (3):309–348.

Felcher, K.J., J.J. Coombs, A.N. Massa, C.N. Hansey, J.P. Hamilton et al., 2012 Integration of Two Diploid Potato Linkage Maps with the Potato Genome Sequence. PLoS ONE 7 (4):e36347.

Fulladolsa, A.C., F.M. Navarro, R. Kota, K. Severson, J.P. Palta et al., 2015 Application of Marker Assisted Selection for Potato virus Y Resistance in the University of Wisconsin Potato Breeding Program. American Journal of Potato Research 92 (3):444–450.

Gebhardt, C., and J.P.T. Valkonen, 2001 Organization of genes controlling disease resistance in the potato genome. Annual Review of Phytopathology 39 (1):79– 102.

Gray, S., S. De Boer, J. Lorenzen, A. Karasev, J. Whitworth et al., 2010 Potato virus Y: An Evolving Concern for Potato Crops in the United States and Canada. Plant Disease 94 (12):1384–1397.

Hackett, C.A., B. Boskamp, A. Vogogias, K.F. Preedy, and I. Milne, 2017 TetraploidSNPMap: Software for Linkage Analysis and QTL Mapping in Autotetraploid Populations Using SNP Dosage Data. J Hered 00 (00):1–5.

Hackett, C.A., J.E. Bradshaw, and G.J. Bryan, 2014 QTL mapping in autotetraploids using SNP dosage information. Theoretical and Applied Genetics 127 (9):1885–1904.

Hackett, C.A., J.E. Bradshaw, and J.W. McNicol, 2001 Interval Mapping of Quantitative Trait Loci in Autotetraploid Species. Genetics 159 (4):1819–1832.

Hackett, C.A., K. McLean, and G.J. Bryan, 2013 Linkage Analysis and QTL Mapping Using SNP Dosage Data in a Tetraploid Potato Mapping Population. PLoS ONE 8 (5):e63939.

Hamilton, J., C. Hansey, B. Whitty, K. Stoffel, A. Massa et al., 2011 Single nucleotide polymorphism discovery in elite north american potato germplasm. BMC Genomics 12 (1):302.

Harrell Jr., F.E., 2016 Hmisc: Harrell Miscellaneous. R package version 4.0-0 https://cran.r-project.org/package=Hmisc.

Hirsch, C.N., C.D. Hirsch, K. Felcher, J. Coombs, D. Zarka et al., 2013 Retrospective View of North American Potato (Solanum tuberosum L.) Breeding in the 20th and 21st Centuries. G3: Genes/Genomes/Genetics 3 (6):1003–1013.

Hosaka, K., Y. Hosaka, M. Mori, T. Maida, and H. Matsunaga, 2001 Detection of a simplex RAPD marker linked to resistance to Potato virus Y in a tetraploid potato. American Journal of Potato Research 78 (3):191–196.

Jones, R.A.C., 1990 Strain group specific and virus specific hypersensitive reactions to infection with potyviruses in potato cultivars. Annals of Applied Biology 117 (1):93–105.

Karasev, A., and S. Gray, 2013a Continuous and Emerging Challenges of Potato virus Y in Potato. Annual Review of Phytopathology 51:571-586.

Karasev, A.V., and S.M. Gray, 2013b Genetic Diversity of Potato virus Y Complex. American Journal of Potato Research 90 (1):7–13.

Kasai, K., Y. Morikawa, V.A. Sorri, J.P. Valkonen, C. Gebhardt et al., 2000 Development of SCAR markers to the PVY resistance gene Ryadg based on a common feature of plant disease resistance genes. Genome 43 (1):1–8.

Kerlan, C., and B. Moury, 2008 Potato virus Y, pp. 287-296 in Encyclopedia of Virology (Third Edition), edited by B.W.J. Mahy and M.H.V.v. Regenmortel. Academic Press, Oxford.

Li, X., Y. Wei, A. Acharya, Q. Jiang, J. Kang et al., 2014 A Saturated Genetic Linkage Map of Autotetraploid Alfalfa (Medicago sativa L.) Developed Using Genotyping-by-Sequencing Is Highly Syntenous with the Medicago truncatula Genome. G3: Genes/Genomes/Genetics 4 (10):1971–1979.

Massa, A.N., N.C. Manrique-Carpintero, J.J. Coombs, D.G. Zarka, A.E. Boone et al., 2015 Genetic Linkage Mapping of Economically Important Traits in Cultivated Tetraploid Potato (Solanum tuberosum L.). G3: Genes/Genomes/Genetics 5 (11):2357–2364.

McDonald, J.G., and R.P. Singh, 1996 Response of potato cultivars to North American isolates of PVYNTN. American Potato Journal 73 (7):317–323.

Michelmore, R.W., and B.C. Meyers, 1998 Clusters of Resistance Genes in Plants Evolve by Divergent Selection and a Birth-and-Death Process. Genome Research 8 (11):1113–1130.

Moury, B., B. Caromel, E. Johansen, V. Simon, L. Chauvin et al., 2011 The Helper Component Proteinase Cistron of Potato virus Y Induces Hypersensitivity and Resistance in Potato Genotypes Carrying Dominant Resistance Genes on Chromosome IV. Molecular Plant-Microbe Interactions 24 (7):787–797.

Ottoman, R.J., D.C. Hane, C.R. Brown, S. Yilma, S.R. James et al., 2009 Validation and Implementation of Marker-Assisted Selection (MAS) for PVY Resistance (Ry_adg_ gene) in a Tetraploid Potato Breeding Program. American Journal of Potato Research 86 (4):304–314.

PGSC, 2011 The Potato Genome Sequencing Consortium. Genome sequence and analysis of the tuber crop potato. Nature 475 (7355):189–195.

Preedy, K.F., and C.A. Hackett, 2016 A rapid marker ordering approach for high-density genetic linkage maps in experimental autotetraploid populations using multidimensional scaling. Theoretical and Applied Genetics 129 (11):2117–2132.

R Core Team, 2016 A Language and Environment for Statistical Computing. R Foundation for Statistical Computing, Vienna, Austria.

Rezvoy, C., D. Charif, L. Guéguen, and G.A.B. Marais, 2007 MareyMap: an R-based tool with graphical interface for estimating recombination rates. Bioinformatics 23 (16):2188–2189.

Sato, M., K. Nishikawa, K. Komura, and K. Hosaka, 2006 Potato virus Y Resistance Gene, Ry_chc_, Mapped to the Distal End of Potato Chromosome 9. Euphytica 149 (3):367– 372.

Schubert, J., V. Fomitcheva, and J. Sztangret-Wiśniewska, 2007 Differentiation of Potato virus Y strains using improved sets of diagnostic PCR-primers. Journal of Virological Methods 140 (1–2):66–74.

Schwarz, G., 1978 Estimating the Dimension of a Model. The Annals of Statistics 6 (2):461–464.

Sharma, S.K., D. Bolser, J. de Boer, M. Sønderkær, W. Amoros et al., 2013 Construction of Reference Chromosome-Scale Pseudomolecules for Potato: Integrating the Potato Genome with Genetic and Physical Maps. G3: Genes/Genomes/Genetics 3 (11):2031–2047.

Singh, R.P., M. Singh, and J.G. McDonald, 1998 Screening by a 3-primer PCR of North American PVYN isolates for European-type members of the tuber necrosis-inducing PVYNTN subgroup. Canadian Journal of Plant Pathology 20 (3):227–233.

Solomon-Blackburn, R.M., and H. Barker, 2001 A review of host major-gene resistance to potato viruses X, Y, A and V in potato: genes, genetics and mapped locations. Heredity 86 (1):8–16.

Song, Y.-S., L. Hepting, G. Schweizer, L. Hartl, G. Wenzel et al., 2005 Mapping of extreme resistance to PVY (Rysto) on chromosome XII using anther-culture-derived primary dihaploid potato lines. Theoretical and Applied Genetics 111 (5):879–887.

Song, Y.-S., and A. Schwarzfischer, 2008 Development of STS Markers for Selection of Extreme Resistance (Rysto) to PVY and Maternal Pedigree Analysis of Extremely Resistant Cultivars. American Journal of Potato Research 85 (2):159–170.

Sorri, V.A., K.N. Watanabe, and J.P.T. Valkonen, 1999 Predicted kinase-3a motif of a resistance gene analogue as a unique marker for virus resistance. Theoretical and Applied Genetics 99 (1):164–170.

Szajko, K., M. Chrzanowska, K. Witek, D. trzelczyk-Żyta, H. Zagórska et al., 2008 The novel gene Ny-1 on potato chromosome IX confers hypersensitive resistance to Potato virus Y and is an alternative to Ry genes in potato breeding for PVY resistance. Theoretical and Applied Genetics 116 (2):297–303.

Szajko, K., D. Strzelczyk-Żyta, and W. Marczewski, 2014 Ny-1 and Ny-2 genes conferring hypersensitive response to Potato virus Y (PVY) in cultivated potatoes: mapping and marker-assisted selection validation for PVY resistance in potato breeding. Molecular Breeding 34 (1):267–271.

Valkonen, J., K. Wiegmann, J. Hämäläinen, W. Marczewski, and K. Watanabe, 2008 Evidence for utility of the same PCR-based markers for selection of extreme resistance to Potato virus Y controlled by Rysto of Solanum stoloniferum derived from different sources. Annals of Applied Biology 152 (1):121–130.

Voorrips, R.E., 2002 MapChart: Software for the Graphical Presentation of Linkage Maps and QTLs. Journal of Heredity 93 (1):77–78.

Watanabe, K., 2015 Potato genetics, genomics, and applications. Breeding Science 65 (1):53–68.

Wei, T., and V. Simko, 2016 Corrplot: Visualization of a Correlation Matrix. R package version 0.77. https://cran.r-project.org/package=corrplot.

Xu, Y., and J.H. Crouch, 2008 Marker-Assisted Selection in Plant Breeding: From Publications to Practice. Crop Science 48 (2):391–407.

